# Fast and Accurate Distance-based Phylogenetic Placement using Divide and Conquer

**DOI:** 10.1101/2021.02.14.431150

**Authors:** Metin Balaban, Yueyu Jiang, Daniel Roush, Qiyun Zhu, Siavash Mirarab

**Affiliations:** Bioinformatics and Systems Biology Graduate Program, University of California San Diego, CA 92093, USA; Center for Fundamental and Applied Microbiomics, Arizona State University, Tempe, AZ 85281, USA; Department of Electrical and Computer Engineering, UC San Diego, CA 92093, USA

**Keywords:** Phylogenetic placement, Distance-based methods, Metagenomics, Microbiome

## Abstract

Phylogenetic placement of query samples on an existing phylogeny is increasingly used in molecular ecology, including sample identification and microbiome environmental sampling. As the size of available reference trees used in these analyses continues to grow, there is a growing need for methods that place sequences on ultra-large trees with high accuracy. Distance-based placement methods have recently emerged as a path to provide such scalability while allowing flexibility to analyze both assembled and unassembled environmental samples. In this paper, we introduce a distance-based phylogenetic placement method, APPLES-2, that is more accurate and scalable than existing distance-based methods and even some of the leading maximum likelihood methods. This scalability is owed to a divide-and-conquer technique that limits distance calculation and phylogenetic placement to parts of the tree most relevant to each query. The increased scalability and accuracy enables us to study the effectiveness of APPLES-2 for placing microbial genomes on a data set of 10,575 microbial species using subsets of 381 marker genes. APPLES-2 has very high accuracy in this setting, placing 97% of query genomes within three branches of the optimal position in the species tree using 50 marker genes. Our proof of concept results show that APPLES-2 can quickly place metagenomic scaffolds on ultra-large backbone trees with high accuracy as long as a scaffold includes tens of marker genes. These results pave the path for a more scalable and widespread use of distance-based placement in various areas of molecular ecology.

## 1 Introduction

Phylogenetic placement of query samples on an existing phylogeny is increasingly used in diverse downstream applications such as microbiome profiling (Asnicar et al., 2020; Darling et al., 2014; Janssen et al., 2018; Matsen, 2014; Matsen and Evans, 2013; Nguyen et al., 2014; Thompson et al., 2017), genome skimming (Bohmann et al., 2020), and epidemic tracking (Libin et al., 2017; Turakhia et al., 2020). A main attraction of placing new sequences onto an existing phylogeny is computational expediency: the running time of phylogenetic placement is a fraction of the time needed for *de novo* reconstruction and can grow linearly with the number query samples assuming they are placed independently. To take advantage of this potential, many methods have been developed using a wide range of algorithmic techniques (e.g., Balaban and Mirarab, 2020; Barbera et al., 2019; Brown and Truszkowski, 2013; Jiang et al., 2021; Linard et al., 2019; Matsen et al., 2010; Mirarab et al., 2011; Rabiee and Mirarab, 2018; Stark et al., 2010; Zheng et al., 2018).

A major attraction of phylogenetic placement is that it enables placement of sequences on very large trees. In applications of placement for microbiome analyses, sequences obtained from amplicon sequencing or metagenomic samples are placed into a reference phylogeny composed of known organisms. Depending on the datatype and pipeline, we may decide to place reads directly (Mirarab et al., 2011; Nguyen et al., 2014) or may place marker genes obtained from metagenome-assembled genomes (MAGs) (Asnicar et al., 2020). Large 16S databases have existed for more than a decade (DeSantis et al., 2006; Quast et al., 2012) and genome-wide references trees with ten thousand species and more have been developed recently (e.g., Parks et al., 2020; Zhu et al., 2019). Moreover, close to a million microbial genomes are available in the RefSeq and GenBank databases. Although there is much redundancy among assembled genomes, we can expect that even larger and more diverse reference trees will be available in the near future. Development of bigger reference sets has a strong motivation: the density of reference set has been known to play a crucial role in the accuracy of downstream analyses (McDonald et al., 2015; Nayfach et al., 2019; Pasolli et al., 2019). Thus, if downstream methods can handle them, we should ideally use these dense reference data sets.

Despite their promise, two types of challenges emerge when reference data sets increase in size: scalability and accuracy. The issue of scalability is well-understood: placement methods may not be able to place on ultra-large reference trees with reasonable running time, and equally important, with reasonable amounts of memory. Moreover, handling ultra large reference trees can be subject to numerical issues. Less appreciated is the observation that as the data set size increases, the accuracy of the algorithms may reduce and updated strategies may be needed. Thus, for placement methods to reach their full potential and take advantage of the available ultra-large reference trees, both scalability and accuracy of existing methods need to improve.

One recent advance in phylogenetic placement on ultra-large reference trees is the development of distance-based placement method, implemented in a method called APPLES (Balaban et al., 2020). Distance-based placement relies on computing distances between the query and references and finding the placement most congruent with these distances. In extensive simulation studies, Balaban et al. (2020) found APPLES to come very close in accuracy to a leading maximum likelihood method, pplacer (Matsen et al., 2010), but unlike ML methods, was able to scale to trees with up to 200,000 taxa. Moreover, APPLES is more useful for studying ecological data because it allows assembly-free and alignment-free placement of genome skims. Despite the relatively high accuracy and scalability of APPLES, it has room for improvement. Its memory usage and speed both grow linearly with the size of the data set, which can start to become slow for references with many hundreds of thousands of species. A bigger challenge is that computing distances across very diverse species found in ultra-large trees can lead to low accuracy, an issue that APPLES only tried to address using weighted distances. A more direct algorithm that accounts for very diverse sequences in the backbone has the potential to further improve accuracy. Moreover, APPLES lacked several features that help usability (including handling of amino acids and building precomputed reference packages). Finally, APPLES has not been tested in the context of microbiome analyses with large backbone trees where it has the most potential.

In this paper, we introduce APPLES-2, a method that, compared to APPLES, improves both accuracy and scalability by adding a divide-and-conquer mechanism and several other features. We test APPLES-2 on both simulated and empirical data sets representative of microbiome analyses. We show that it can place scaffolds from a metagenomic sample onto a large reference tree of more than 10,000 species both given marker genes found in assembled scaffolds.

## 2 Materials and Methods

### 2.1 APPLES-2 Algorithm

#### 2.1.1 Background

Balaban et al. (2020) introduced a Least Squares Phylogenetic Placement (LSPP) framework and a method called APPLES for distance-based placement. In this framework, the input to APPLES is a reference (*a*.*k*.*a* backbone) phylogenetic tree *T* with *n* leaves and a vector of distances *δ*_*qi*_ between a query taxon *q* and every taxon *i* on *T*. Although machine-learning based methods shows substantial promise (Jiang et al., 2021), typically, input distances are obtained by calculating sequence distances between query and backbone taxa followed by a phylogenetic correction using a statistically consistent method under a model such as Jukes-Cantor (JC69) (Jukes and Cantor, 1969). APPLES introduced a dynamic programming algorithm to find a placement of *q* that minimizes weighted least squares error 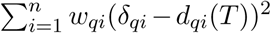 where *d*_*qi*_(*T*) represents the path distance from *q* to backbone taxon *i* on *T*. APPLES, by default, sets 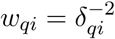 following Fitch and Margoliash (1967) (FM) weighting.

#### 2.1.2 Divide-and-conquer Placement Algorithm

The most consequential change in APPLES-2 is that it adopts a divide-and-conquer approach to improve both accuracy and scalability using two inter-related techniques. There is strong evidence in distance-based phylogenetics literature that correction for high variance occurring in estimation of long distances can result in dramatic improvements in accuracy (Desper and Gascuel, 2002; Felsenstein, 2003; Whitfield, 2008). For example, the DCM family of methods that result in fast converging methods (Erdos et al., 1999; Huson et al., 1999a,b) mostly rely on dividing taxa into smaller subsets with lower distances. To take advantage of this insight, we enable APPLES-2 to use distances that are either smaller than a threshold *d*_*f*_ or among the lowest *b* distances. Ignoring distances larger than the *d*_*f*_ threshold also gives us an opportunity to avoid computing all *n* distances so that the running time could grow sublinearly with the size of the reference tree. To do so, we divide the backbone tree *T* into subsets that are somewhat larger than *d*_*f*_ in diameter (maximum pairwise path distance between any two leaves), choose one representative from each subset, and compute distances of the query only to these representatives. Then, we compute all distances in the cluster with the least distance to our query taxon.

More formally, without loss of generality, we assume that for a certain query taxa *q, δ*_*q*1_ ≤ *δ*_*q*2_ ≤ *δ*_*q*3_ ≤ … *δ*_*qn*_ holds. The first parameter we introduce is *d*_*f*_ ∈ ℝ_*≥*0_ which sets *w*_*qi*_ = 0 (i.e. ignore backbone taxon *i*) when *δ*_*qi*_ ≥ *d*_*f*_ for a query taxon *q*. In addition, we introduce a second parameter *b* ∈ ℕ_*≥*0_ which forces to retain the standard weighting (i.e.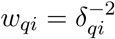) for backbone taxa 1 ≤ *i* ≤ *b*, regardless of *δ*_*qi*_. Consequently the new LSPP objective function becomes 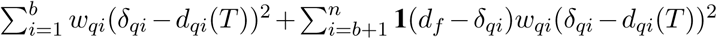 where **1**(*x*) is the unit step function: **1**(*x*) = 0 for *x <* 0 and **1**(*x*) = 1 for *x* ≥ 0. We discuss default values below.

To avoid computing all distances, during preprocessing of the reference set, we cluster the backbone alignment and tree *T* using the linear-time TreeCluster algorithm (Balaban et al., 2019) to find the minimum number of clusters such that the maximum pairwise distance in each cluster is no more than 1.2 × *d*_*f*_. The threshold 1.2 is chosen empirically, and APPLES-2 is robust to this choice (see Fig. S1). Then, we select a representative sequence per partition by computing consensus sequence among all sequences belonging to the partition. Let *P*_1_, *P*_2_, …, *P*_*k*_ denote partitions of leaves of *T* and *C*(*j*) denote centroid sequence of partition *P*_*j*_. Without loss of generality, we assume that *δ*_*qC*(1)_ ≤ *δ*_*qC*(3)_ ≤ *δ*_*qC*(3)_ ≤ … *δ*_*qC*(*k*)_. The distance between *q* and a backbone taxa *i* ∈ *P*_*j*_ is computed only if either *δ*_*qC(j)*_ ≤ *d*_*f*_ or 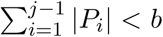 holds. The time complexity of distance calculation per query is in the order of *O*(max(*b, m*)*L*) where *L* is alignment length and *m* is number of backbone taxa whose distance to the query is less than or equal to *d*_*f*_.

Since in APPLES-2 a subset of distances are calculated, we have re-designed its dynamic programming algorithm so that it automatically works on the backbone tree induced to the taxa for which distances are computed. The updated dynamic programming algorithm scales with the number of edges in the induced tree, which can be as low as *O*(max(*b, m*)) (if the chosen leaves are a connected subtree) and as high as *O*(*n*) (when chosen leaves span all of a caterpillar tree).

#### 2.1.3 New features in APPLES-2 Software

##### Protein distances

Several tools (Lefort et al., 2015; Rice et al., 2000; Sonnhammer and Hollich, 2005; Womble, 2000) offer distance calculation from protein sequences using analytical (Jukes and Cantor, 1969; Kimura, 1983; Sonnhammer and Hollich, 2005) and maximum likelihood (ML) (Le and Gascuel, 2008; Whelan and Goldman, 2001) models. In order to provide support for protein alignments, we implement Scoredist algorithm, which has achieved better accuracy than other analytical models in previous tests (Sonnhammer and Hollich, 2005). Scoredist computes pairwise distances according to the BLOSUM62 (Henikoff and Henikoff, 1992) matrix, normalizes the distances with respect to expected distance and minimum possible distance, applies a logarithmic correction, and scales distances using empirically derived coefficients. Like JC69 distances, in APPLES-2, Scoredist distance calculation is powered by Numpy (Harris et al., 2020) vectorized operations and is extremely fast.

##### BME weighting

APPLES implemented three weighting schemes FM (Fitch and Margoliash, 1967), BE (Beyer et al., 1974), and OLS (Cavalli-Sforza and Edwards, 1967). Balaban et al. (2020) demonstrated that FM weighing given by 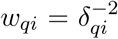 results in the best placement accuracy among these methods. However, it did not implement balanced minimum evolution (BME) weighting (Desper and Gascuel, 2004), which has been among the most promising methods. APPLES-2 implements BME which corresponds to setting 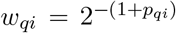 where *p*_*qi*_ is the topological distance between *q* and a backbone taxa *i*. Note that BME weights are much more challenging to incorporate into the dynamic programming because a BME weight is not simply a function of calculated distances but is rather a function of the placement on the tree. Thus, unlike the previous weighting schemes, the BME weight changes as we examine different placements. Overcoming this hurdle required implementing a more complex dynamic programming.

##### Database features

We allow precomputation of a database (called APPLES database) that consists of a backbone alignment and tree, including centroid sequences and the leaf clustering, which can be stored and distributed. The database can be reused for the analysis of different query data sets. Moreover, when a backbone alignment is provided, APPLES-2 can re-estimate branch lengths of the input tree using FastTree-2 (Price et al., 2010) under the JC69 model to match the model used for estimating distances.

### 2.2 Experiments

#### 2.2.1 Data sets

##### RNASim data set

We reuse the RNASim-VS simulated RNA data set from Balaban et al. (2020), which consists of subsets of a simulated RNASim data set (Guo et al., 2009), but change the query selection strategy. We begin with randomly selecting 200 queries with various novelty levels; to control novelty, 10 taxa are randomly selected from each of 20 bins determined by dividing terminal branch length of the phylogeny on 200,000 taxa into 20 quantiles. The remaining 199,800 taxa are designated as backbone. Then, we create data sets with size (*n*): 100000, 50000, 10000, 5000, 1000, and 500 by successive random subsampling. The procedure is replicated 5 times and query taxa are identical within a replicate across different size data sets. Each replicate contains a 1596-sites multiple-sequence alignment of a single gene and the true tree. 200 query are placed on the backbone independently for each replicate. We also adopted the RNASim-QS data set from Balaban et al. (2020) also based on the RNASim data set (Guo et al., 2009); this data set comprises five replicates with varying numbers of queries, ranging from 1 to 49,152 with backbones of size *n* = 500. In both RNASim data sets, backbone tree topology and maximum likelihood branch lengths are estimated from true MSA using FastTree-2 (Price et al., 2010) according to GTR+Γ model. In all cases, branch length is re-estimated using FastTree-2 (Price et al., 2010) to be consistent minimum evolution units.

##### Web of Life (WoL) data set

Zhu et al. (2019) built a species tree of 10,575 prokaryotic genomes from 381 marker genes using ASTRAL (Zhang et al., 2018). We first determine a set of marker genes according to several selection strategies which will be discussed later. We remove sites that contain gaps in 95% or more of the sequences in the protein MSA using TrimAl (Capella-Gutierrez et al., 2009). The trimming is only to speed up analyses and has no positive impact on accuracy; in fact, it very slightly *decreases* accuracy (see Fig. S2). Then, we create three concatenated alignments; the amino acid alignment, a nucleotide alignment with all three codon positions (C123), and another with third codon position removed (C12). Unless it is stated otherwise, we use the C12 nucleotide MSA in our analyses.

We analyze the WoL data set in four ways (Table. 1). In WoL-main, three data sets of size (*n*) 9000, 3000, and 1000 with 10 replicates are created by successively subsampling the protein MSA of the selected marker genes at random. From the remaining 1575 species, 1000 are randomly subsampled from the protein MSA of the selected marker genes and designated as query. For all data set sizes, we use the ASTRAL tree available from the original publication induced to backbone species as the backbone tree. However, we recompute its branch lengths using FastTree-2 (Price et al., 2010) in the minimum evolution branch length unit. We determine a marker gene set by controlling for two parameters; the number of genes (*k*) and a selection strategy. Two selection strategies are *random* (among all 381) and *best*, which means top *k* marker genes with the lowest quartet distance (Sand et al., 2013) to the species tree are selected. In WoL-main, we choose *k* = 50 coupled with the *best* strategy (which results in lowest, median, and highest quartet-distance to be 0.058, 0.125, 0.17 respectively). Concatenated MSA using the default marker gene set contains 71, 798 nucleotide sites. In WoL-random (Table 1), we create 1000-species backbone alignments by selecting *k* ∈ {10, 25, 50, 381} coupled with the *best* and *random* strategies. Additionally, marker gene set selection is replicated five times for the *random* strategy. In the previous two data sets, the backbone was inferred with queries included, which were then removed, because repeating the complex backbone inference pipeline for all analyses was not doable. However, we did add a smaller analysis that avoids this information leakage. In WoL-denovo, we reuse a single replicate under data set sizes 1000 and 3000 from WoL-main and fully reproduce WoL pipeline (Zhu et al., 2019) to obtain de-novo MSA and species tree instead of removing queries from the full tree. All query genes are then independently aligned to de-novo MSA of the 50 backbone marker genes using UPP (Nguyen et al., 2015).

**Table 1:** WoL based data sets. In Wol-random data set, only random selection of marker genes is replicated 5 times. *best* marker selection strategy indicates choosing the marker genes whose gene tree has the lowest topological discordance with the species tree. An alignment or backbone tree is induced when it is taken from a larger data set (e.g. full data set). C12: nucleotide alignment with first and second codon positions. C123: nucleotide alignment with all codon positions. AA: amino-acid alignment.

| data set name | Backbone Size | Number of markers | Marker strategy | Backbone tree & MSA | Query alignment | Replicates | Character |
| --- | --- | --- | --- | --- | --- | --- | --- |
| WoL-main | 1000, 3000, 9000 | 50 | best | induced | induced | 10 | C12, C123, AA |
| WoL-random | 1000 | 10, 25, 50, 381 | best, random | induced | induced | 5* | C12 |
| WoL-denovo | 1000, 3000 | 50 | best | de-novo | UPP | 1 | C12 |
| WoL-metagenomic | 10375 | 381 | all | induced | UPP | 1 | C12 |

##### Data set of Simulated genome assemblies and scaffolds

In WoL-metagenomic data set, we utilize a protocol for generating simulated genome sequencing data which begins with randomly selecting 200 test genomes from the WoL data set (10 genomes are randomly selected from each of 20 genome bins of equal genome count with the bins determined by ascending terminal branch length). Next, we run InSilicoSeq (Gourlé et al., 2019) v1.5.1 (using NovaSeq settings) to simulate 3 M 150 bp paired-end reads per genome. For assembly, first we run PEAR (Zhang et al., 2014) v0.9.11 to merged read pairs, then run SPAdes (Bankevich et al., 2012) v3.14.1 with a *k*-mer size cascade of 21,33,55,77,99 to assemble them into scaffolds. We then run Prodigal (Hyatt et al., 2010) v2.6.3 to identify open reading frames (ORFs) from the scaffolds, and finally run PhyloPhlAn (Segata et al., 2013) commit 2c0e61a to identify the same 381 marker genes.

Selected test genomes are removed from the backbone set, which leaves us with 10375 species in the backbone. All the genes were then independently aligned to the backbone marker genes using UPP (Nguyen et al., 2015), and markers from the same assembly or scaffold were concatenated. We try to place the samples on the backbone using either 1) the assembly (i.e., which can be fragmented and can include errors, compared with the genome from which it is simulated), or 2) individual scaffolds (small portions of the genome). We only include scaffolds that are ≥ 10 kbp in our analyses. Note that here, instead of testing on microbial communities, we use an *in-silico* approach and simply generate reads from individual microbial genomes and assemble them separately. We leave it to future work to simulate mixed metagenomic reads and evaluate accuracy under such scenarios.

##### Biological TD-metagenomic dataset

We use the metagenome-assembled genomes (MAGs) from a study by Zhu et al. (2018), which identified novel pathogenic profiles from the fecal samples of 22 Traveler’s Diarrhea (TD) patients and seven healthy traveler (HT) controls. The dataset consists of 320 manually curated MAGs (bins) and 6653 scaffolds that are 50kb or longer. The 381 marker genes in the dataset are identified using the same protocol as the WoL study. We use the species tree in WoL study as the backbone tree and align the sequences from Traveler’s Diarrhea dataset to the WoL dataset using UPP independently for each marker gene. We then concatenate the 381 marker genes from the same bins or scaffolds and use them for placement. We also study the case where we filter out the scaffolds with less than or equal to 10, 20, 30, or 40 marker genes, which reduce the number to 4522, 1608, 668, and 320 scaffolds, respectively.

#### 2.2.2 Methods compared

For APPLES-2, we explored various options for *d*_*f*_ and *b* in an experiment performed on the WoL-main data set (Fig. S3). As a result, we set *d*_*f*_ = 0.2 and *b* = 25 by default and keep these values fixed across all of our other experiments. For RNASim-VS and WoL-main data set, in addition to APPLES, we compare APPLES-2 to two ML methods, pplacer (Matsen et al., 2010) and EPA-ng (Barbera et al., 2019). We run pplacer (v1.1.alpha19-0-g807f6f3) and EPA-ng (v0.3.8) in their default mode using GTR+Γ model and use their best hit (ML placement). Unlike the procedure used by Balaban et al. (2020), we do not perform branch length re-estimation on backbone tree using RAxML-8 (Stamatakis, 2014). Instead, we input inferred FastTree-2 tree and model parameters without modification to EPA-NG and pplacer. We compare to EPA-ng in analyses that concerned scalability (e.g., RNASim-QS).

#### 2.2.3 Evaluation Criteria

In RNASim analyses, we use the known true tree as the gold standard whereas on the empirical data, we use the ASTRAL tree on the full set of species as the gold standard with an exception of WoL-denovo data set in which the ASTRAL tree is computed denovo for each data set size. In all WoL data sets except WoL-denovo, we measure the accuracy of a placement using the number of edges between the position on the gold-standard tree and the inferred placement (i.e., node distance (Linard et al., 2020)). In the simulated RNAsim data set, because true tree is known and we place on the estimated tree and not the true tree, we need to slightly update the metric of the error: We use *delta error*, which measures the increase in the number of false-negative bipartitions after placement compared to before placement (Mirarab et al., 2011). We use delta error in WoL-denovo data set as well, treating the published phylogeny on the full set as the true tree.

## 3 Results

### 3.1 Single-gene Placement

We start with leave-one-out experiments on an existing single-gene simulated RNASim data set where all methods face model misspecification. Despite the model misspecification, APPLES-2 is able to find the best placement of query sequences with up to 91% accuracy (placement on the correct branch) when the backbone size is *n* = 200, 000 (Fig.1a, Table 2). APPLES-2 has lower mean delta error (−0.08 edges on average) and higher accuracy (+2.5% on average) compared to APPLES for all cases except for *n* ≤ 5000, where they are essentially tied in accuracy but APPLES has slightly higher mean delta error. Across all cases, pplacer is the most accurate method. In particular, pplacer has 10% better accuracy and 0.13 less mean delta error than APPLES-2 for *n* = 500. However, the difference in accuracy and mean error gradually decrease as *n* increases and diminish to only 2% and 0.04 respectively for *n* = 200, 000. Compared to the other ML method, EPA-NG, APPLES-2 either matches (for *n* ≤ 5000) or improves the accuracy (up to 3%) on instances where EPA-NG manages to complete (*n* ≤ 100, 000). Thus, APPLES-2 matches or improves the accuracy of one ML method (EPA-ng) and is slightly below the accuracy of the other (pplacer).

**Figure 1:**
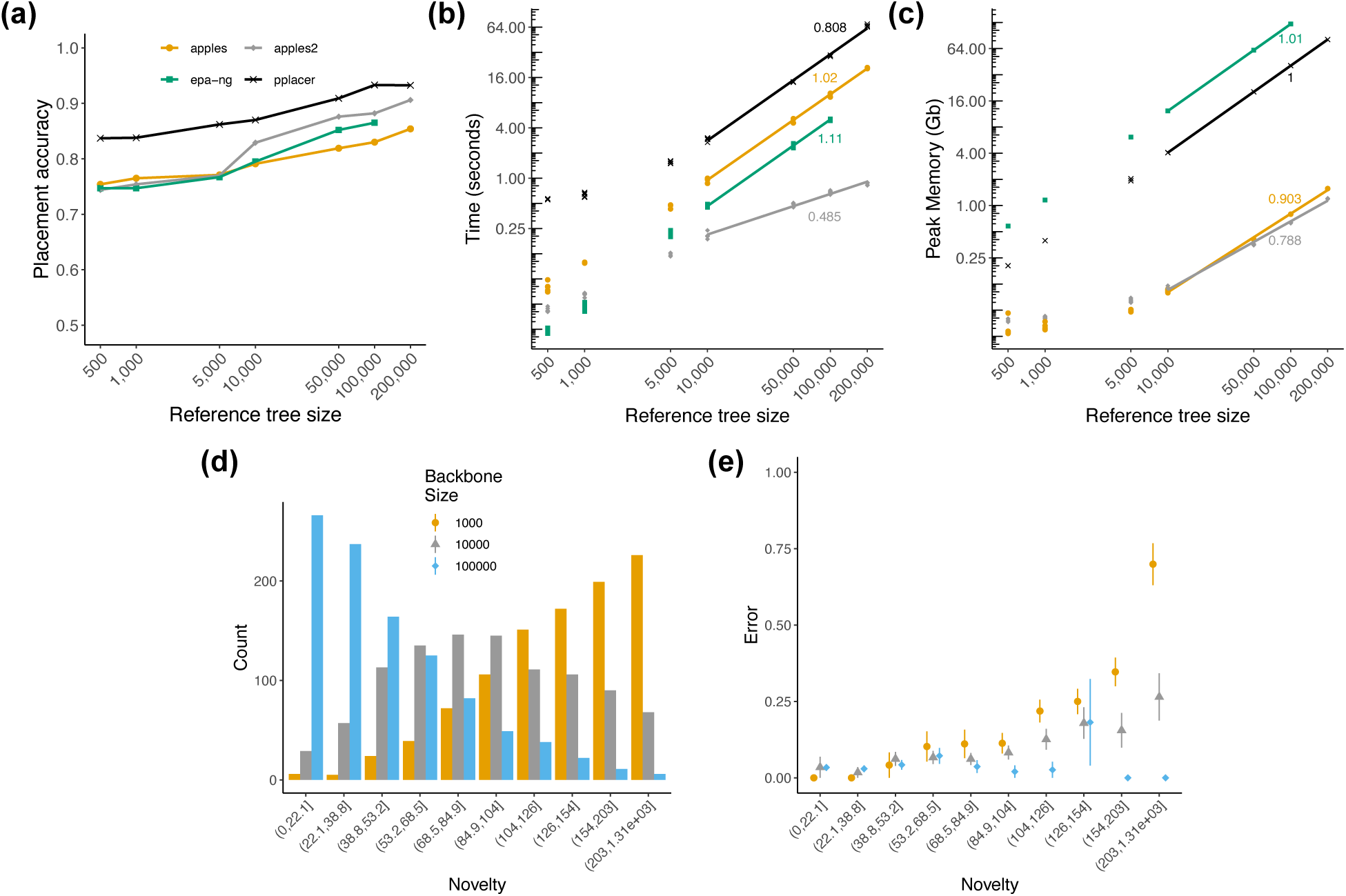
Results on RNASim-VS. Placement accuracy (a), running time (b), and peak memory usage (c) for a single placement with taxon sampling ranging from 500 to 200,000. (b,c) Lines are fitted in the log–log scale and their slope (indicated on the figure) empirically estimates the polynomial degree of the asymptotic growth. Lines are fitted to ≥10000 points because the earlier values are small and irrelevant to asymptotic behavior. All calculations are on 36-core, 2.6GHz Intel Xeon CPUs (Sandy Bridge) with 128GB of memory, with each query placed independently and given 1 CPU core and the entire memory. Missing results (EPA-NG on tree size 200, 000) indicate that the tool fails to run or complete in 48h. (d) Queries are grouped into deciles based on their novelty with respect to backbone set of species, defined as the terminal branch length of the query in the gold-standard tree, induced to backbone and query species. (e) Mean placement error of APPLES-2 across increasing level of query novelty for all three backbone sizes.

**Table 2:** Percentage of correct placements (shown as %) and the average placement error (Δ*e*) on the RNASim-VS with various backbone size (*n*). % and Δ*e* is over 1000 placements (except *n* = 200, 000, which is over 200 placements). *n*.*p* indicates tool failed to run in the case.

|  | n = 500 |  | n = 1,000 |  | n = 5,000 |  | n = 10000 |  | n = 100,000 |  | n = 200,000 |  |
| --- | --- | --- | --- | --- | --- | --- | --- | --- | --- | --- | --- | --- |
| | % | $\Delta e$ | % | $\Delta e$ | % | $\Delta e$ | % | $\Delta e$ | % | $\Delta e$ | % | $\Delta e$ |
| **APPLES-2** | 74 | 0.31 | 75 | 0.33 | 77 | 0.34 | 83 | 0.24 | 88 | 0.14 | 91 | 0.11 |
| **APPLES** | 75 | 0.36 | 77 | 0.34 | 77 | 0.36 | 79 | 0.34 | 83 | 0.28 | 85 | 0.22 |
| **EPA-ng** | 75 | 0.29 | 75 | 0.28 | 77 | 0.24 | 80 | 0.21 | 87 | 0.14 | n.p | n.p |
| **pplacer** | 84 | 0.18 | 84 | 0.18 | 86 | 0.14 | 87 | 0.14 | 93 | 0.07 | 93 | 0.07 |

Placement accuracy of APPLES-2 is 17% higher on largest tree than the smallest tree. To examine the reason, we first observe that the novelty of the test set (defined as terminal branch length in the true tree) decreases as the backbone size increases (Fig 1D). To test the impact of novelty on error, we measure the mean error for each decile of novelty for all backbone sizes after larger trees are pruned so that backbone trees are identical to those of the smallest tree. This pruning ensures that errors are always computed with respect to trees of the same size and are therefore comparable. Two patterns stand out. First, increasing novelty does increase the error, especially for smaller backbone sizes (Fig 1E). Thus, the error with larger backbone trees is reduced simply because fewer queries are novel (Fig 1D). More interestingly, it appears that at higher levels of novelty, the error is reduced with larger backbones even after the novelty level is controlled. Thus, the results show improved accuracy with the increased taxon sampling even when the novelty of the test set does not change. We note that larger backbone trees include fewer long branches in the backbone and that processes such as long branch attraction need at least two close long branches (e.g., one in the backbone and one for the query) to impact results.

Our benchmarking indicates that the running time of APPLES-2 grows empirically as *O*(*n*^0.45^) (Fig. 1b); this sublinear running time growth with the backbone size is consistent with our theoretical expectations. APPLES-2 is the fastest method on backbones with 5,000 or more taxa, offering up to 24× speed-up in average compared to APPLES on a 200,000-taxon tree. EPA-NG is faster than pplacer and APPLES but slower than APPLES-2 (with running time that grows super-linearly). On the 100,000 taxon data set, EPA-NG and pplacer take 7.3× and 77× longer than APPLES-2 on average respectively. APPLES-2 and APPLES consistently uses less memory than ML tools (Fig. 1c) and are the only tools with sublinear memory complexity (empirically close to *O*(*n*^0.8^) for APPLES-2). On the largest backbone tree with 200, 000 taxa, APPLES-2 requires only 1.2GB of memory compared to 81GB needed by pplacer. EPA-NG uses 192× more memory then APPLES-2 on the largest backbone tree with 100, 000 taxa where both tools successfully run.

We also evaluated the impact of the number of queries on the running time (Fig. 2), comparing APPLES, APPLES-2, and EPA-NG, all run in the parallel mode. On 500 taxa backbone, all three methods finish placement of up to 1,536 queries in less than 4 seconds given 28 CPU cores with no clear trend in running time. EPA-NG is able to place 49152 queries in 10 seconds on average, 5.8 times faster than the second best method APPLES-2, which takes 57 seconds and is 6.5 times faster than APPLES. The comparison of EPA-NG and APPLES-2, the fastest two of the three methods, on backbone trees with 1000 and 5000 taxa shows that EPA-NG is 6 and 3.4 faster than APPLES-2 respectively on the largest query set. While both methods complete in less than 36 seconds, APPLES-2 is faster than EPA-NG when number of queries is less than or equal to 1536 for 5000 taxa tree. Running times of EPA-NG, which is designed specifically for very large numbers of query sequences, can surprisingly decrease when given more queries. For any backbone size, APPLES and APPLES-2 start to have substantial increase in running time after placing 6144 queries or, scaling linearly with respect to the number of queries; surprisingly, EPA-NG grows at a sublinear rate, likely indicating that it requires more queries to display its asymptotic behavior. To summarize, while APPLES-2 is faster than EPA-NG given hundreds of queries, EPA-NG scales better as the number of queries increases.

**Figure 2:**
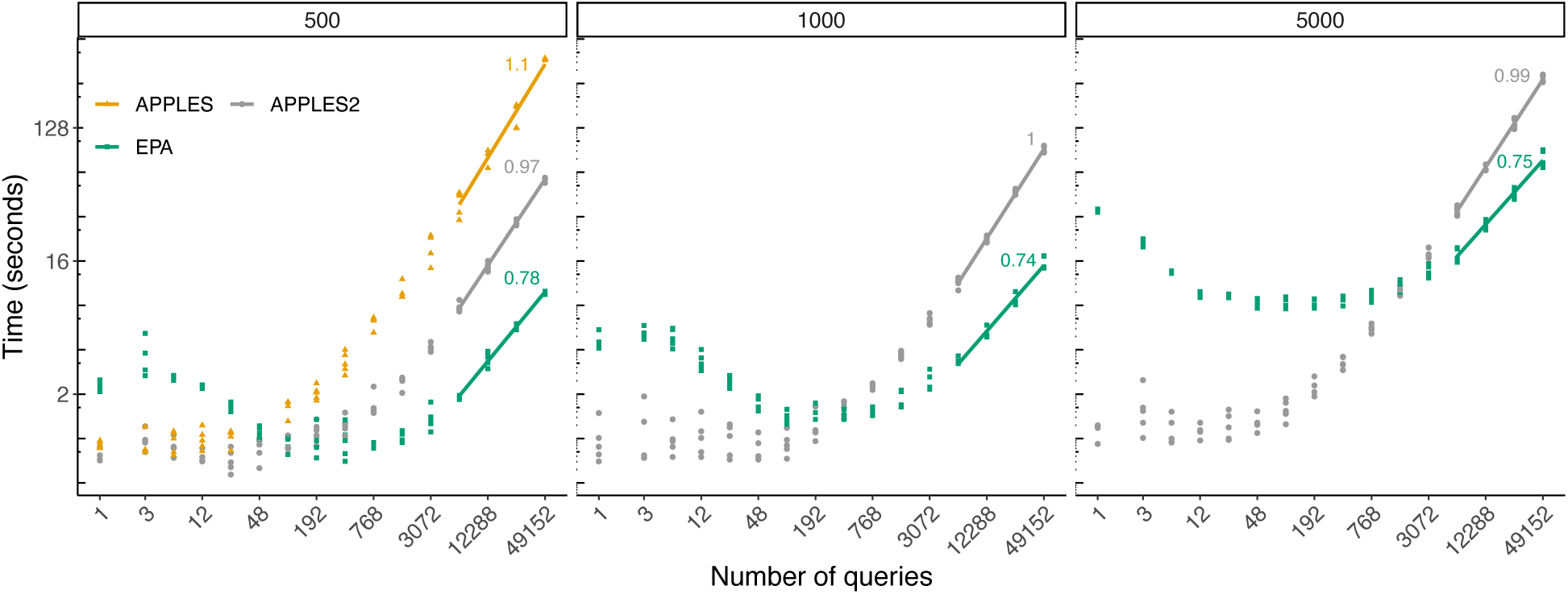
Scalability with respect to the number of queries. We show wall-clock running time with respect to increased numbers of queries placing on a tree with 500 taxa in one execution of the tool given 28 CPU cores and 28 threads on an Intel Xeon E5 CPU with 64 GB of memory. Lines are fitted to *x* ≥ 6144 points because the earlier values are small and irrelevant to asymptotic behavior.

### 3.2 Multi-gene Web of Life (WoL) data set

We next test the utility of distance-based phylogenetic placement on a real WoL biological data set (Zhu et al., 2019) with 381 marker genes and 10575 microbial taxa. When we concatenate the best 50 marker genes, APPLES-2 achieves outstanding accuracy, placing query sequences with 75% accuracy and 0.50 edges of error on average on backbones with 1000 taxa (Fig. 3a). A striking 97% of the queries are placed within three or fewer branches away from the optimal branch (in a tree with a diameter of 58.3 branches on average). Note that here we are using 50/381 marker genes and a much simpler methodology compared to the original study. In comparison, APPLES achieves 60% accuracy with 1.1 average error on the same data set. As in the single-gene RNASim dataset, pplacer is the most accurate method with 80% accuracy for *n* = 1000. EPA-NG has slightly lower accuracy (−1%) and mean error (−0.06) than APPLES-2. As the size of the reference increases from 1000 to 3000 and 9000, APPLES-2 and APPLES are only methods that run successfully due to large memory requirements of ML-based methods (more on performance below). APPLES-2 is able to maintain high accuracy, placing within three branches of the optimal placement in 97% and 96% of cases, respectively, for backbones of size 3000 (avg. diameter: 77.2 edges in average) and 9000 (avg. diameter: 105.5 edges). Increasing the backbone size also amplifies the gap between APPLES and APPLES-2, going from a difference of 0.57 edges of error on average for *n* = 1000 to 0.81 and 1.11 for *n* = 3000 and *n* = 9000.

**Figure 3:**
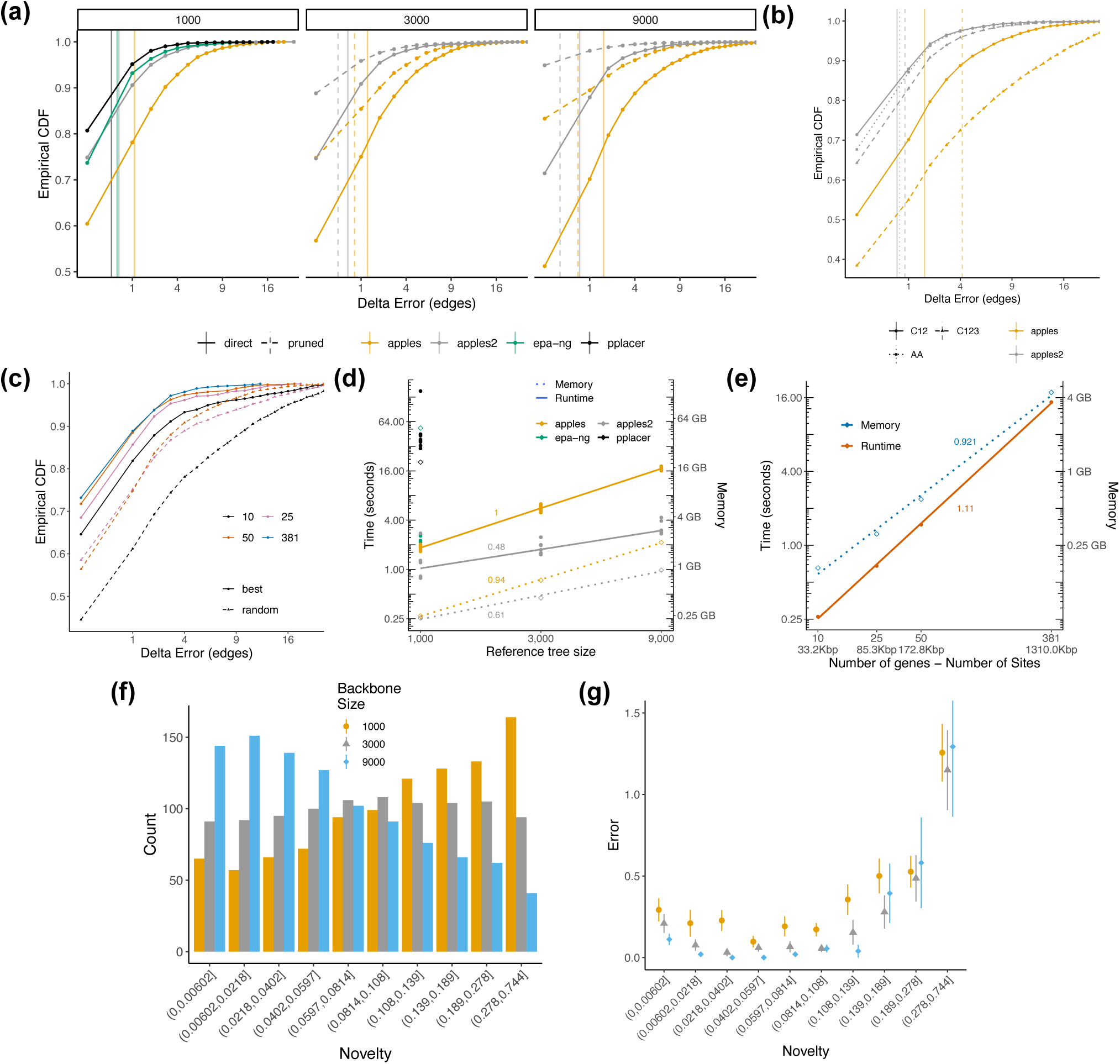
(a) Empirical cumulative distribution function (CDF) of placement error on backbones ranging from 1000 to 9000 taxa. Results are based on 10000 query placement for each backbone size. Pruned delta error is calculated after pruning the placement tree to species in the *n* = 1000 tree and queries. Vertical lines show mean error. X-axis is displayed in square-root scale and is cut at 20 edges. (b) Placement accuracy versus alignment type. C12: only the first two codon positions are retained in the alignment; C123: all three positions used. (c) Impact of marker gene selection on placement accuracy. We control for number of genes selected and gene selection strategy: choosing randomly versus genes with lowest discordance with species tree (best). (d,e) Running time (solid lines and solid points) and memory (dotted lines, hollow points) performance with respect to backbone tree size (d) and number of marker genes in the backbone tree (e). Lines are fitted in the log–log scale and their slope empirically estimates the polynomial degree of the asymptotic growth. Each run has 32 cores and 56GB memory in a shared node with 2.25 GHz AMD EPYC 7742 processor with each query placed independently and given 1 CPU core and the entire allocated memory. Missing results indicate that the tool fails to run or complete in 48h. (f,g) Novelty (defined as in Fig. 1) of queries and mean placement error of APPLES-2 for all backbone sizes.

APPLES-2 places queries with 0.1 higher error in average for *n* = 9000 compared to *n* = 1000; however, it should be noted that the largest tree has 9 times more branches than the smallest one. Therefore one branch of error in the smallest tree indicates a larger degree of misplacement. In order to establish a fair comparison between trees with different number of backbone species, after placement (i.e. before measuring the error), we prune trees with *n* = 3000 and *n* = 9000 to include only those present in the smallest tree with *n* = 1000. The comparisons on pruned trees shows that the placement accuracy for APPLES-2 becomes 90% and 95% on 3000 and 9000-taxa backbone tree respectively, which are much higher than 75% with 1000-taxa backbone tree (Fig. 3a). These increases in accuracy show that accuracy of APPLES-2, does, in fact, improve with better taxon sampling.

The reasons behind improved accuracy with improved taxon sampling parallel the simulated data. Again, we observe reduced novelty in the query set (Fig. 3f) as backbone size increases. Impact of novelty on error is not uniform. When the query is extremely similar to backbone species, correct placement is difficult. Thus, initially, the error slightly decreases as novelty increases. However, after reaching a sweet spot, the error increases dramatically as novelty increases. The better accuracy with larger trees, therefore, is a function of having fewer very novel queries. Controlling for novelty of query, in the first seven deciles, results show a negative correlation between the error and backbone size (Fig. 3g).

In WoL-main data set, backbone and query alignment and backbone tree is directly induced from full WoL data set, which may potentially “leak” information about query location since query sequences were present in the full data set during MSA and tree inference. We test this scenario on WoL-denovo data set where two MSAs and trees with *n* = 1000 and *n* = 3000 are de-novo inferred using the identical methodology described in the original publication (Zhu et al., 2019). In addition, query sequences are aligned to backbone MSA using UPP (Nguyen et al., 2015) to prevent leakage of information through alignment. We find a slight absolute reduction (−5%) in placement accuracy of APPLES-2 on 3000-taxa de-novo backbone compared to induced backbone (Fig. S4). However, the percentage of queries placed with no more than three edges error is 98% for both de-novo and induced backbone trees. The mean delta experiences very slight changes between de-novo and induced backbone trees.

APPLES-2 is the fastest method in WoL-main data set, managing to place a query in 1.1 second on average on the smallest backbone tree using a single CPU core (Fig. 3d). For comparison, APPLES, EPA-NG, and pplacer take 1.86, 2.18, and 49.47 seconds per query respectively on the same data set. APPLES and APPLES-2 achieve the best memory efficiency by using 250Mb of memory whereas EPA-NG and pplacer use 48.6 and 18.6 GB of memory on the same instances. As backbone size increase to *n* = 3000 and *n* = 9000, APPLES and APPLES-2 become the only methods that complete the benchmark given a 56GB memory machine as ML-based methods terminate due to insufficient memory. Our benchmark indicates that running time and memory use of APPLES-2 grow sublinearly, achieving empirical time and memory complexity of *O*(*n*^0.5^) and *O*(*n*^0.6^) respectively.

Next, testing the impact of data type used for placement, we observe that removing the third codon position from nucleotide alignments improves placement accuracy substantially for both versions of APPLES (Fig. 3b). Interestingly, APPLES-2 seems to be more robust to inclusion of third codon position as the increase in the average error is 0.26 and 2.44 for APPLES-2 and APPLES, respectively. The third codon position often poses a stronger violation on stationarity assumption than the first and second codon positions (Jeffroy et al., 2006; Phillips et al., 2004) and saturates faster, especially among very divergent taxa. Recall that APPLES-2 ignores distances among very divergent sequences, which is consistent with its higher robustness to the third codon position. Note that the original study (Zhu et al., 2019) that built our gold-standard in these analyses inferred gene trees using amino acid data. We do not observe a substantial error difference between using nucleotide (first two codon positions) and amino acid sequences (Fig. 3b) for APPLES-2. Although APPLES-2 has 3% higher accuracy on the former data type, the number of queries with at most three edges of error is 96% on both data types. We remind the reader that amino acid distances are computed under the Scoredist algorithm, which is different from the models used in the original study to infer the reference tree (Zhu et al., 2019).

Next, we test the impact of varying the number of marker genes and the type of genes used (randomly chosen or the *best* genes) on WoL-random data set (Fig. 3c). While Using all the marker genes has the highest accuracy (mean edge error: 0.52; placement accuracy: 73%), using as few as 50 of best genes (i.e., those with gene trees with the lowest quartet distance the species tree) comes very close. With 50 genes, APPLES-2 places 958 out of 1000 queries (96% rate) within three branches away from the optimal branch; in contrast, using all genes, 972 queries are within three branches (97% rate). Using the best 50 genes results in 0.63 average delta error, which is 0.11 more than using all genes in the data set. However, reducing the number of best genes to 25 and 10 increases error to 0.79 and 1.28 edges, respectively. Our benchmark indicates that runtime and memory use of APPLES-2 empirically grow near linearly with number of marker genes and number of sites in the backbone alignment (Fig. 3e). When all 50 marker genes used, placement of a query takes 1.5 seconds whereas using all 381 marker genes, placement takes 10 times longer on the 1000 taxa backbone. Thus, 50 best genes is the sweet spot in terms of accuracy among levels we test considering computational requirements.

There is a large difference between selecting genes randomly or using best genes (Fig. 3c). A random selection of 10 genes results in lower accuracy (within 3 edges from the optimal branch only 74% of the time) and a high average edge error of 3.29, whereas 10 best genes result in 1.28 edges of error on average. Going to 25 randomly selected genes provide acceptable placement accuracy where 87% of queries are placed within 3 edges from the optimal location (error: 1.8 edges on average); yet, 25 best genes continue to be better (error: 0.79 edges on average). With 50 genes, the error is 0.80 less when using best genes compared to randomly selected genes. The mean placement error using random genes decreases as the number of genes increases, culminating in 0.53 edges when all genes are selected (Fig. S5). Note that *best* and *random* strategies are equivalent when all 381 genes in the data set are used. Overall, the difference between random and best genes is wider when few genes are available and diminishes as more genes are added.

### 3.3 Placement of assemblies and scaffolds

While our previous analyses showed that APPLES-2 has outstanding accuracy using best or random subsets of marker genes sampled across microbial genomes, we often do not have entire genomes. Instead, we have MAGs and scaffolds from which MAGs are generated. We next test APPLES-2 in a simulation that generated scaffolds and assemblies, similar to MAGs, by assembling reads simulated from a subset of 200 genomes in the WoL data set.

Our simulated assemblies included 105 to 365 marker genes (Fig. S6a). With these number of markers, APPLES-2 achieves 67% accuracy and places 195 of 200 simulated assemblies with error no larger than three edges (Fig. 4b). The error is never more than 6 edges. The placement error has a weak but statistically significant anti-correlation with the number of marker genes available in the assembly (*p* = 0.002 according to Pearson’s correlation; *ρ* = −0.216; see Fig. S7).

**Figure 4:**
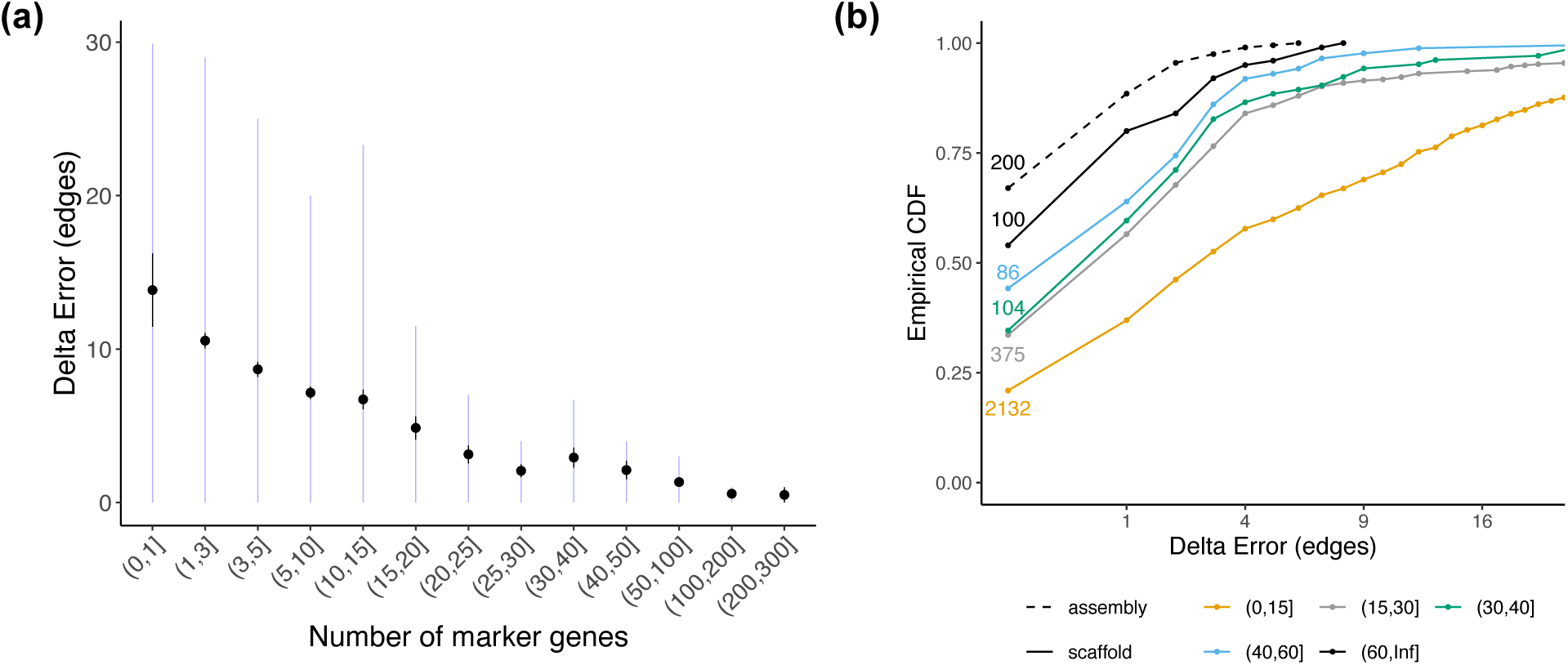
Results on WoL-metagenomic data set. (a) The relationship between number of marker genes in a scaffold and the error. Dots show mean, black error bars show standard error, and light blue error bars show the central 80% range. X-axis is binned non-linearly. (b) Placement error CDF for simulated metagenomic assemblies and scaffolds. Each bin indicates the number of genes in the scaffold or assembly. We show the number of queries in each bin next to its curve. The backbone has the diameter (the largest pairwise distance) of 106 edges.

Our assembly procedure produces 3318 unique scaffolds of ≥ 10kbp (Fig. S6b), among which 665 has more than fifteen marker genes and 290 has more than 30 marker genes. The placement error is clearly a function of the number of genes in each scaffold (Fig. 4a). Scaffolds with less than 15 genes have high error on average (8.51 edges), but also have high variance (with 53% of such scaffolds leading to error up to three edges). Once scaffolds start to have more than approximately 20 genes, the error becomes consistently low (Fig. 4a). The placement accuracy for scaffolds that contains 30 to 40 genes is 35%, and 83% can be placed with error no more than 3 edges (Fig. 4b). As the number of genes in the scaffold increases, the accuracy also increases; on average, placement error for scaffolds with 50 or more genes is only 1.19 edges, 92% are within three edges of the optimal placement, and the maximum error observed is 12 edges.

Both multiple sequence alignment using UPP and the phylogenetic placement step using APPLES-2 used in the scaffold placement workflow are fast. Running UPP to align all 3318 scaffolds for each gene to the backbone alignment takes 89 seconds in average (lowest 18 and highest 388 seconds) using 6 CPU cores. APPLES-2 takes 2.77 seconds per query scaffold on the backbone tree of size of nearly 10000 species using 28 CPU cores.

### 3.4 Placement of real MAGs and scaffolds onto WoL tree

Next, we study the Zhu et al. (2018) metagenomic dataset composed of gut microbiomes of 22 patients with Traveller’s Diarrhea (TD) and 7 healthy controls (HT). For each subject, we obtain six placement profiles by placing MAGs and scaffolds with five marker occupancy thresholds. We compare two profiles by computing weighted Unifrac distance (Lozupone and Knight, 2005). We observe a statistically significant difference between intra-group (HT and HT, TD and TD) and inter-group (TD and HD) distances with MAGs (p-value 0.01 using standard PERMANOVA test) (Fig. 5a). Using all scaffolds (not including MAGs) intra- and inter-group distances between the samples cannot be distinguished with statistical significance (p = 0.099). However, F-statistic increases after filtering out scaffolds with less than or equal to 10 marker genes. Increasing the scaffold filtering threshold to 40 results in a decrease in the F-statistic (Fig. 5a), indicating that a large proportion of the signal in the data is lost due to over-filtering. Using placement, we can also visualize MAG- and Scaffold-informed community structures of all samples using a Principal Coordinates Analysis (PCoA) (Fig. 5b). MAG-informed community structures provide better delineation of communities dominated by *Escherichia coli* compared to Scaffolds with at least 10 marker genes. Thus, our results show that placing MAGs using APPLES-2 enables inference about community structure of metagenomic samples but the utility of scaffolds is less clear.

**Figure 5:**
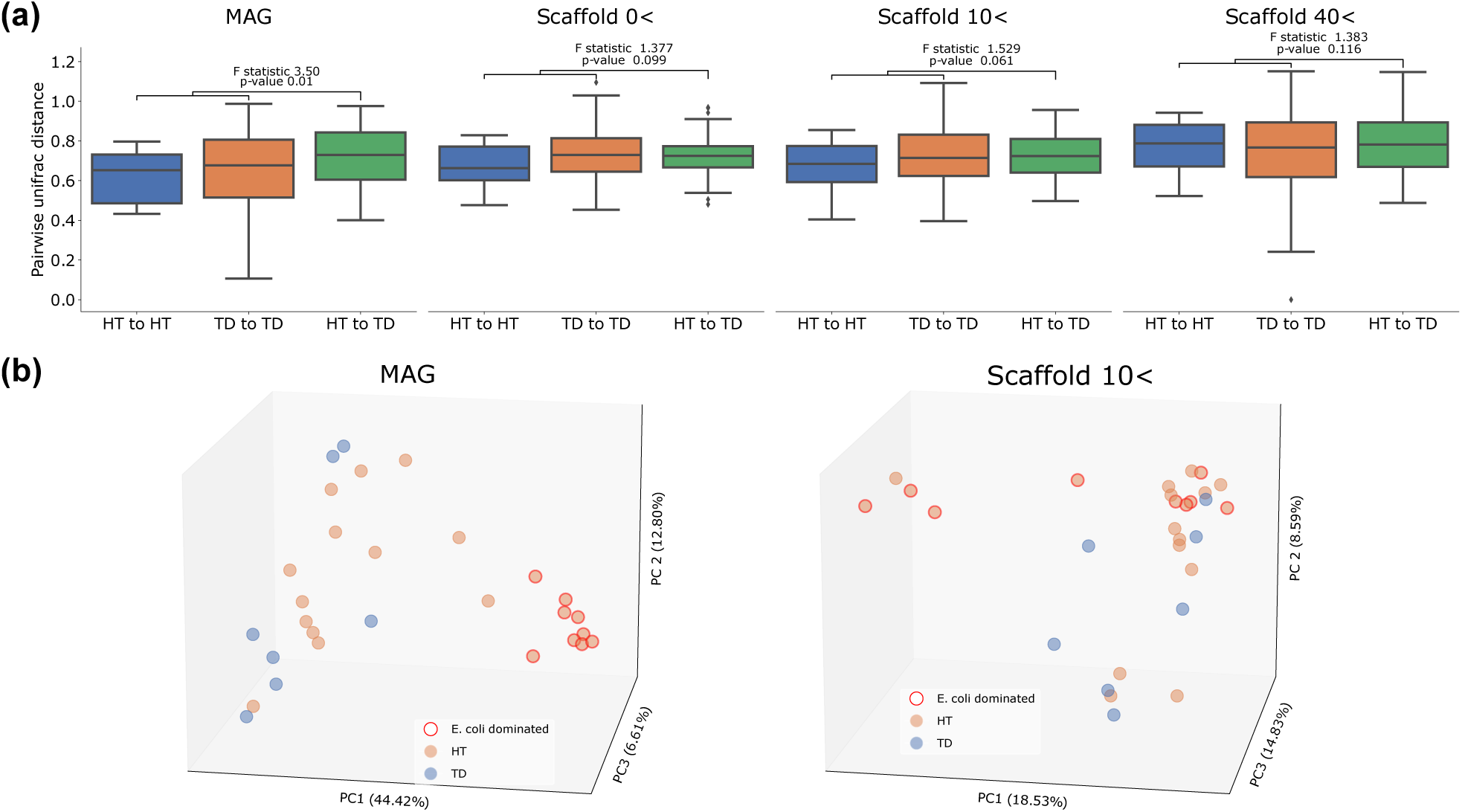
Results on the real TD metagenomic dataset. (a) Distribution of UniFrac distances among pairs of samples within HT or TD group and across the groups, using MAGs and scaffolds with more than 0, 10, or 40 marker genes present. F-statistic and p-values are calculated using PERMANOVA test. (b) The PCoA visualization of microbiome profiles of samples using MAGs and Scaffolds with more than 10 marker genes, highlighting samples known to be dominated by *E. coli*.

## 4 Discussion

We presented APPLES-2: an improved distance based phlogenetic placement tool for inserting new taxa on large phylogenetic trees. Inspired by DCM-like methods (Huson et al., 1999b), our divide-and-conquer approach improved placement accuracy beyond its predecessor APPLES and made it comparable to or better than ML-based tool EPA-NG on single gene data sets. Furthermore, we showed that APPLES-2 is even more scalable than APPLES, reducing running time and memory consumption, and can achieve high accuracy on diverse multi-gene data sets.

Some of the new features of APPLES-2 increase usability and completeness of the tool but have limited impact on accuracy and scalability. For example, we implemented BME weighting. However, despite the previous literature suggesting BME weighting is preferable to alternatives (Desper and Gascuel, 2004), we observed that BME is less accurate than the default FM weighting scheme for all data set sizes (Fig. S8); difference between FM and BME mean error is 0.85 edges on average). Based on these results, we continued to use FM as the default weighting method everywhere but provide BME as a new option to the users. Similarly, using amino acid sequences did not show any improvements, but we enable it for cases when only amino acid data is available. Despite declining opportunities, future changes could seek to further improve the accuracy. For example, at the expense of higher computational cost, one can select centroid sequences for partitions of the backbone MSA via ancestral state reconstruction instead of consensus — a technique used by (Balaban et al., 2019) (also see ancestral k-mers (Linard et al., 2019)).

Previously, Balaban et al. (2020) reported that ML-based method pplacer failed to place queries on backbone trees with 5000 taxa or larger in RNASim-VS data set due to a numerical error (infinity likelihood values). We find that re-estimating backbone branch lengths and model parameters using RAxML-8 and inputting the RAxML info file to pplacer causes a bug in pplacer. We overcome this issue by creating a taxtastic package (https://github.com/fhcrc/taxtastic) using Fasttree-2 tree and info file and using this package as the input. Note that creating taxtastic package from re-estimated RAxML-8 tree and info file also produces the aforementioned error. As a result of discovery, we do not perform branch length re-estimation using RAxML-8 in any of our data sets.

While accuracy is typically high, on a minority of queries results of APPLES-2 are far from the correct placement. A reasonable question is whether these highly inaccurate instances can be identified by APPLES-2. While we leave a more elaborate exploration to future work, we have identified several interesting patterns (Fig. 6). First, we observe a correlation between APPLES-2’s objective function value, the Minimum Least Square Error (MLSE; denoted by *Q*), and placement error (Fig. 6a). In addition, variance of error dramatically increases as *Q* increases. Even for the same level of *Q*, selecting marker genes strategically instead of randomly reduces the placement error. Therefore, *Q* itself does not seem sufficient to predict the degree of placement error. Note that high MLSE (e.g *Q* ≥ 1) does not indicate that APPLES-2 fails to optimize its objective function — APPLES-2 solves the objective problem *exactly* (i.e., is not heuristic). High MLSE can result from sequence data and tree distances being very incompatible. This incompatibility may be due to several reasons such as lack of signal, model violation, and horizontal gene transfer (HGT). Despite its reduced *mean* accuracy, APPLES-2 can still find a good placement for many queries with *Q* ≥ 1: Approximately 75% of such queries have at most 3 edges error on the backbone consisting of random marker genes. Secondly, when a query is placed with zero distal and pendant edge length, the placement error is significantly higher than otherwise (*p <* 5.5 × 10^−13^, two-sample Wilcoxon test). The average error is 11.74 when both pendant and distal edge length is zero (i.e. when query is placed on an internal node) whereas it is only 2.04 on average when pendant edge length is larger than zero (Fig. 6b). We have also noticed that erroneous placements with zero pendant edge lengths are more prevalent in the query sequences with fewer genes. Out of 15 occurrences of this pattern, 13 are found in test cases with 10 marker genes in the backbone. Thus, users of APPLES-2 should be skeptical of the placements with zero pendant and distal branch length and/or high MLSE error (which APPLES-2 outputs). A warning is produced by APPLES-2 when such placement are produced. In future work, these features can be used to develop a predictive value indicating possible errors in placement.

**Figure 6:**
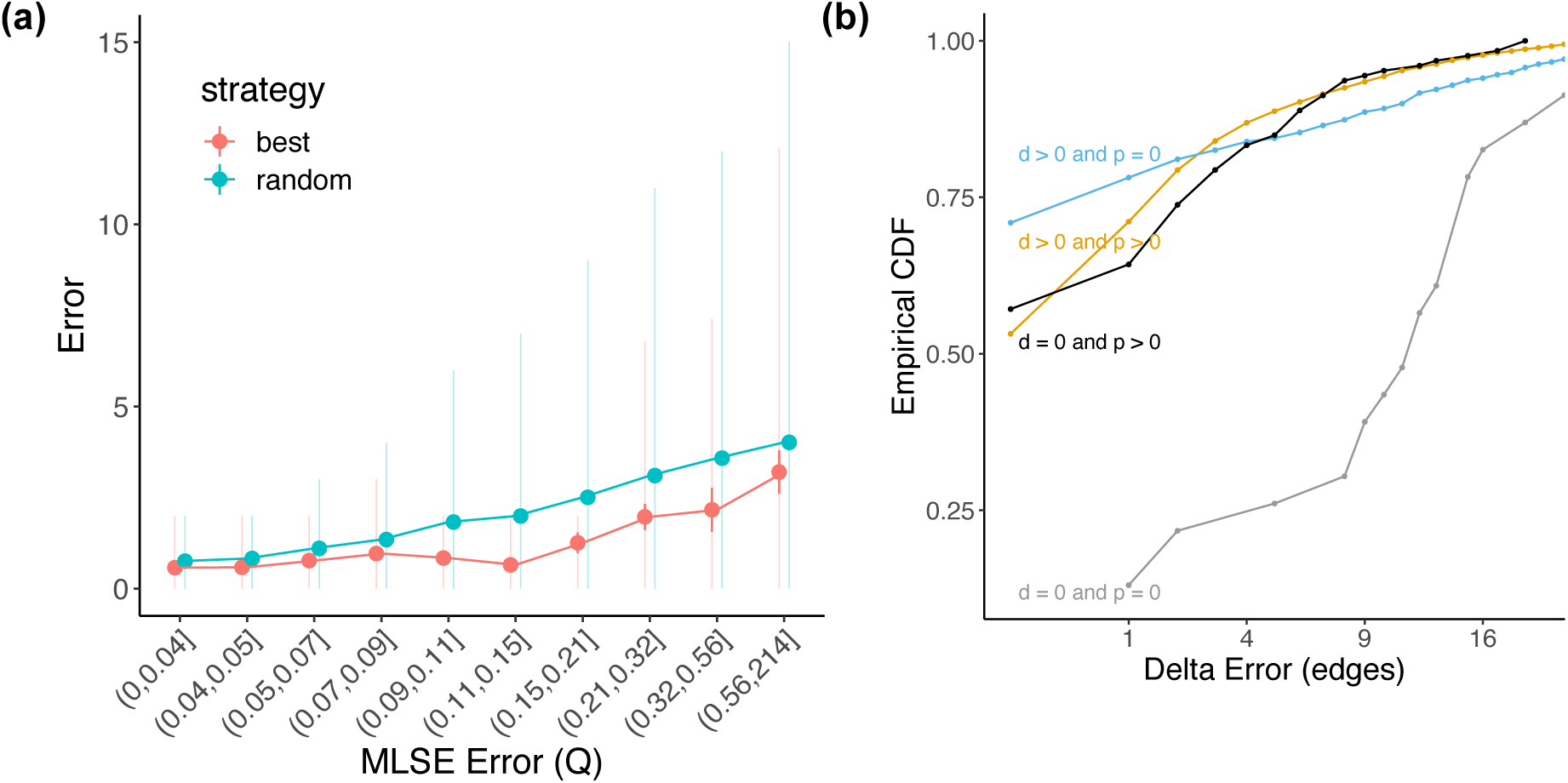
Detecting erroneous placements. Results are based on 18879 queries in WoL-random dataset. (a) The relationship between optimized MLSE objective function (Q) and the error. Dots show mean, thick error bars show standard error, and light error bars show the central 80% range. (b) Empirical CDF of placement error with distal (d) and/or pendant (p) edge equal to zero. Queries with zero objective function (i.e. queries that have an exact match to a backbone species) are omitted.

Our studies on microbial data showed that APPLES-2 can phylogenetically place and hence identify genome-wide shotgun data with promising accuracy after they are assembled. Using both simulated and real data sets, we showed that assembled genomes (MAGs) can be placed on the species tree with great accuracy. Patterns are more intricate for scaffolds: On real data, scaffolds with very few (as few as one) or large number of marker genes (40) were insufficient to portray the community structure of the metagenomic sample. Filtering scaffolds with fewer than ten marker genes provided the optimal signal-to-noise ratio, despite being inferior to MAGs. On simulated microbial data, the accuracy tends to be low on small scaffolds with few genes but improve for scaffolds that have moderately large number of marker genes. Besides their reduced numbers of genes, scaffolds present several challenges that may contribute to their lower accuracy: (*i*) In comparison to assemblies, scaffolds in a metagenomic sample are more prone to assembly errors and chimeras. (*ii*) Genes located on the same syntenic block have similar gene trees, which can introduce a bias in the placement. Thus, factors such HGT and may have a bigger impact on scaffolds. (*iii*) Even when a scaffold has many genes, it may not include the best marker genes; i.e., those genes with maximum signal and concordance to the species tree.

Our results clearly showed that the choice of genes matters. While a random selection of 25 marker genes were adequate for placing queries in most cases, a targeted gene selection strategy outperformed random selection (e.g., *p* = 6.6 × 10^−13^ for 25 marker genes, two-sample Wilcoxon test). The results indicate that certain marker genes serve better at predicting location of a query species in the backbone tree. This observation leads to two related questions. Given fully assembled genomes, should we use all or a subsample of available genes? Our data supports the idea that using a subset of genes has very similar accuracy to using all available genes. However, a more fruitful approach may be weighting genes (or even sites within genes) differently to further improve accuracy. Such a goal seems amenable to machine learning approaches that can learn optimal weights.

The second question is how to handle scaffolds from metagenomic assemblies, which include only a handful of genes. There are always more scaffolds with few genes than those with many genes. Thus, requiring a large number of genes would reduce the number of scaffolds placed, which has the potential to reduce the accuracy of downstream analyses. Our results indicate scaffolds with a modest number of genes (e.g., with 30 or more) are enough to place them phylogenetically. But the vast majority of scaffolds have fewer than 15 marker genes, and *some* of these *can* be placed accurately. We leave it to future work to design a more principled framework for deciding which scaffolds can be placed accurately and which cannot. We also leave to the future work to answer a more challenging question: for downstream applications, is it better to place a few scaffolds that have many genes (or perhaps binned contigs) with high confidence or is it better to place all or most scaffolds with lower confidence hoping that noise will be overcome by the large number of placements? Answering these questions requires careful experimental procedures that are outside the scope of the present study.

While in this paper we focused on applications of APPLES-2 to microbiome data, our earlier work has demonstrated the utility of distance-based placement for assembly-free and alignment-free identification of genome skims (Balaban et al., 2020). While reference sets available for genome skimming are not currently large enough to challenge APPLES in terms of scalability, the divide-and-conquer step in APPLES-2 may lead to increase accuracy. Essentially, the divide-and-conquer mechanism will allow building reference databases that include genome skims from very diverse set of organisms (e.g., all insects) without reducing accuracy due to high levels of divergence. We leave the exploration of such applications and the choice of best thresholds for genome skimming to future work.

Finally, in this paper, we focused on single query placement and observed that given multiple marker genes, APPLES-2 can insert a new genome into backbone tree with high accuracy. These results open up an exciting opportunity. By spending less computational budget than *de novo* phylogenetics, successive insertion of genomes can enable expanding the existing large microbial phylogenies (e.g., Zhu et al., 2019) to contain hundreds of thousands of sequences. Future work should explore the best pipelines for achieving this goal.

## Acknowledgements

We thank Daniel McDonald for his valuable feedback.

## Author Contributions

**MB**: Conceptualization, Methodology, Software, Validation, Formal analysis, Investigation, Data Curation, Writing, Visualization. **YJ**: Formal Analysis, Writing, Visualization. **DR**: Data Curation, Writing. **QZ**: Data Curation, Writing, Supervision. **SM**: Conceptualization, Methodology, Investigation, Writing, Visualization, Supervision.

All authors read and approved the final manuscript.

## Data Availability Statement

The APPLES-2 code is publicly available under GNU GPL-v3 at https://github.com/balabanmetin/apples. The data used in this work are available at https://doi.org/10.5281/zenodo.5551285.

## Appendix A

### Supplementary Figures

**Figure S1:**
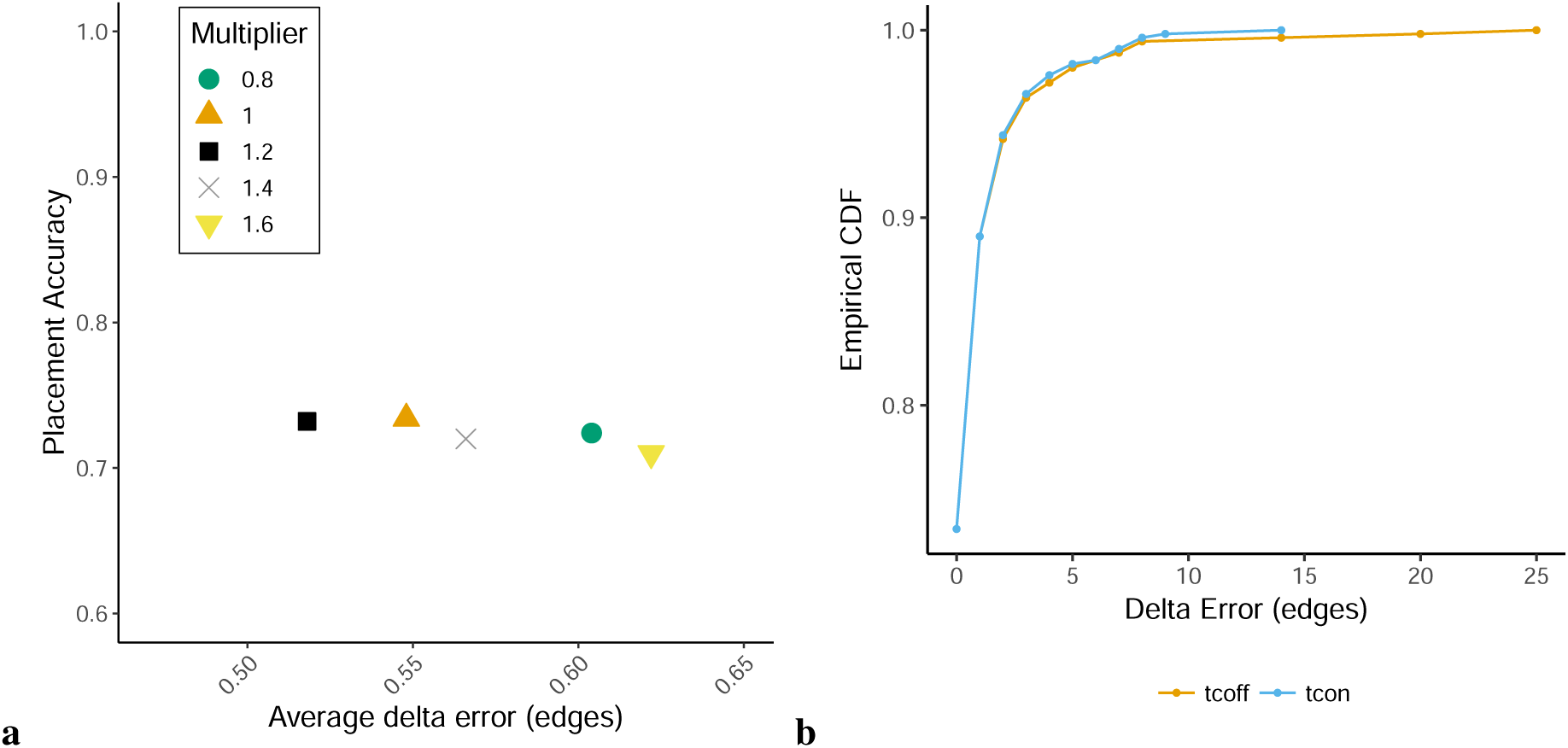
Clustering the backbone using TreeCluster. a) Exploring the multiplier for TreeCluster maximum diameter threshold. Results are based on 500 placements on WoL species tree using APPLES-2. Placement accuracy does not change as this threshold changes and the average error increases slightly if other values are used. b) Using TreeCluster to divide the backbone tree into smaller subsets (shown as *tcon* in the figure) not only improves the running time but also slightly improves the placement accuracy compared to using APPLES-2 on the full backbone tree (shown as *tcoff* in the figure).

**Figure S2:**
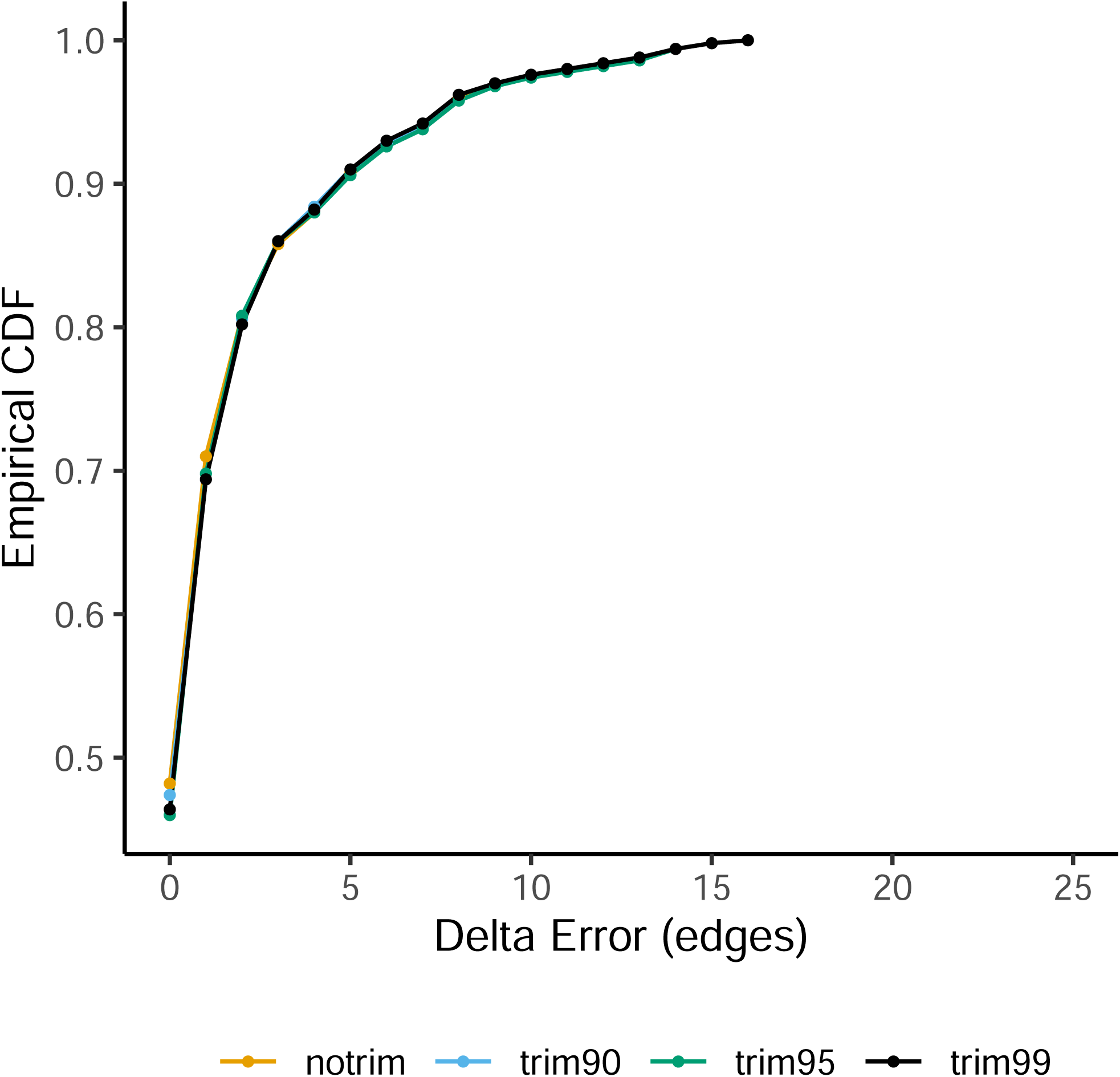
Impact of alignment trimming. Results are based on 500 placements on WoL backbone tree using APPLES. Trimming sites with 90, 95, or 99 percent gaps or more does not dramatically impact placement accuracy compared to using untrimmed alignment. Trimming reduces accuracy only slightly. For improved speed and memory usage, we therefore trim the sites with 95% or more gaps.

**Figure S3:**
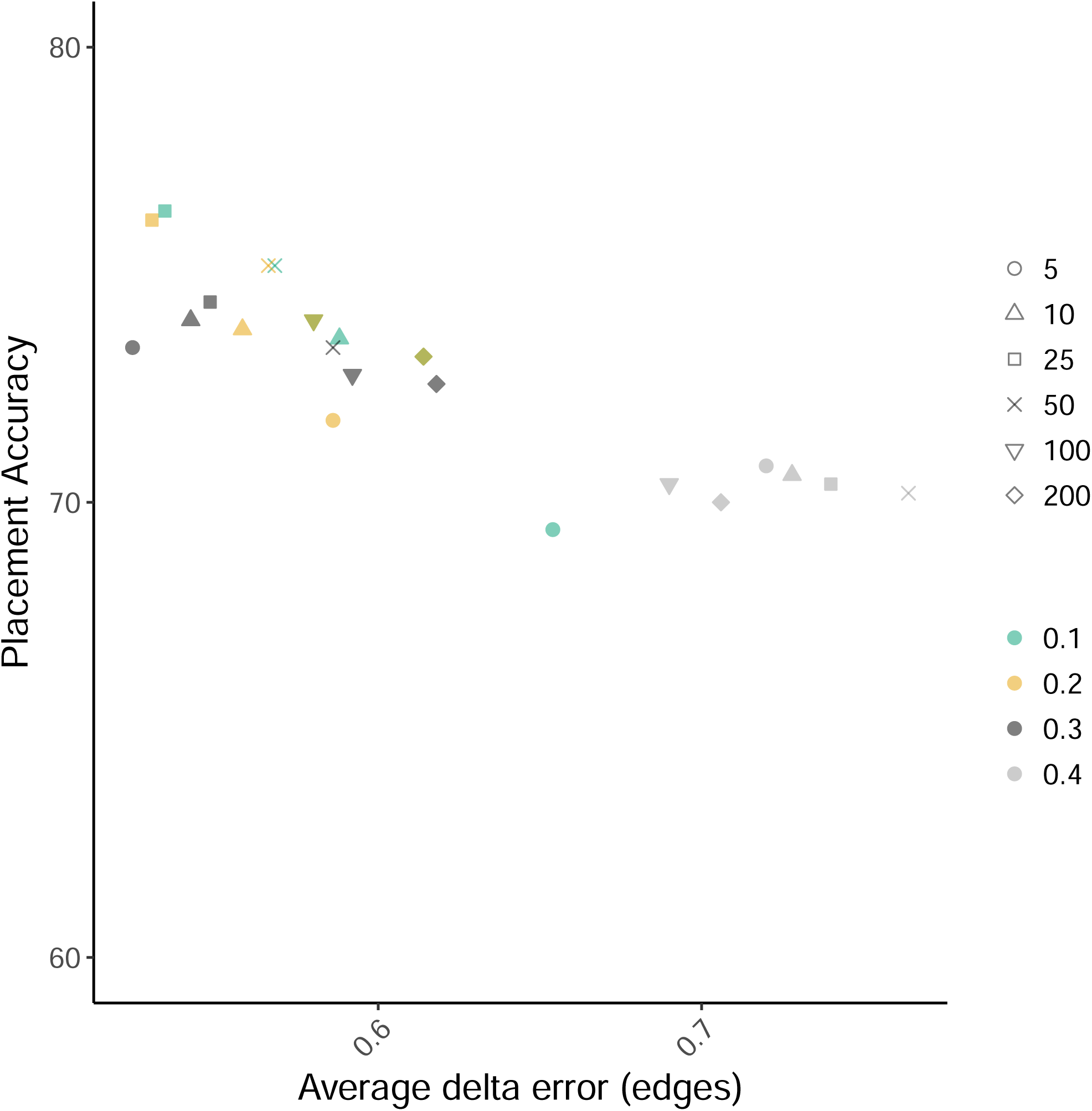
Exploring the best choice of *d*_*f*_ and *b* parameters. Results are based on 500 placements on WoL data set using APPLES-2. Color indicates *d*_*f*_ parameter and shape indicates *b* parameter.

**Figure S4:**
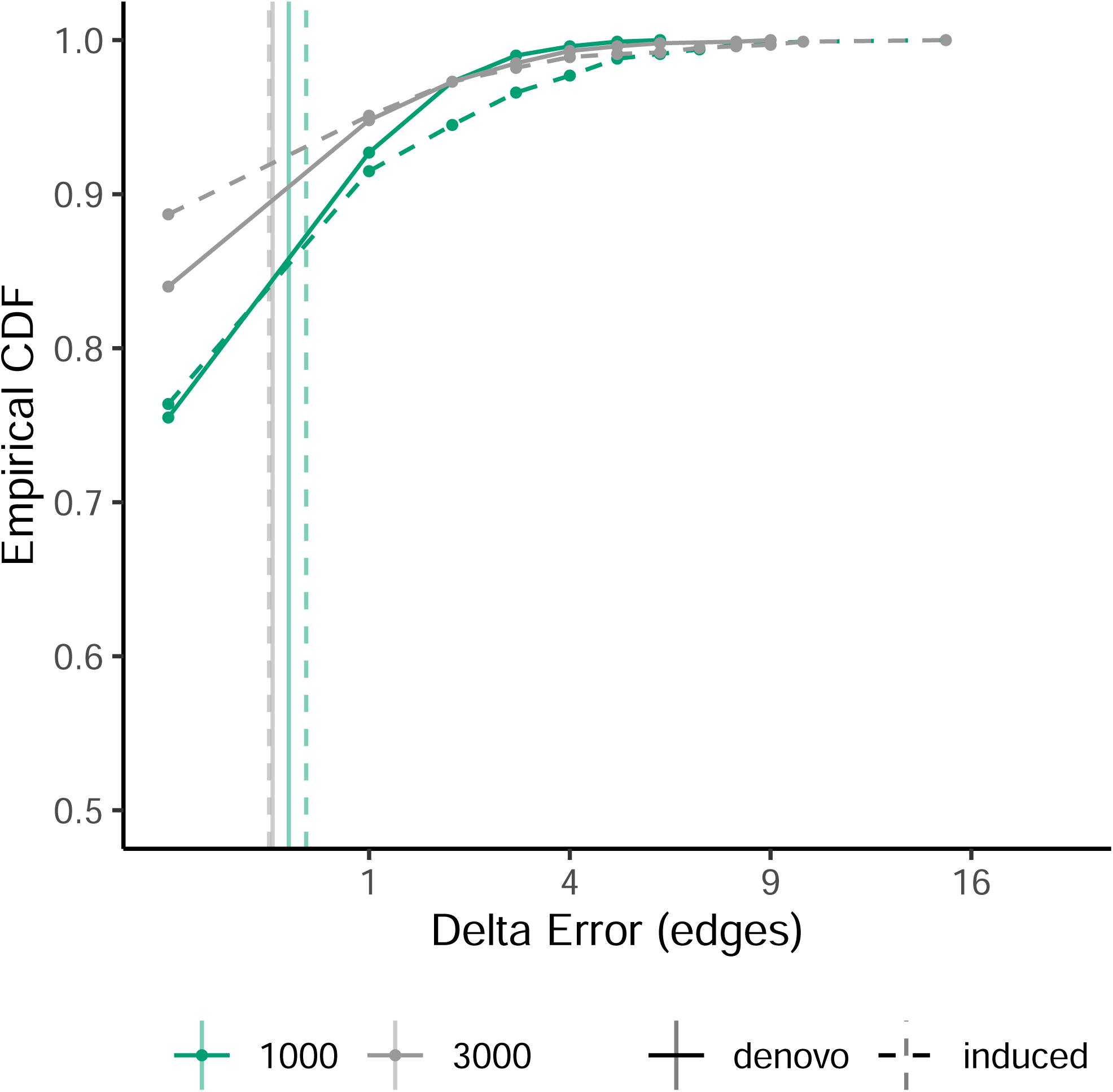
De-novo versus induced backbone. For *n* = 3000, delta error is calculated after pruning the placement tree to species in the *n* = 1000 tree and queries.

**Figure S5:**
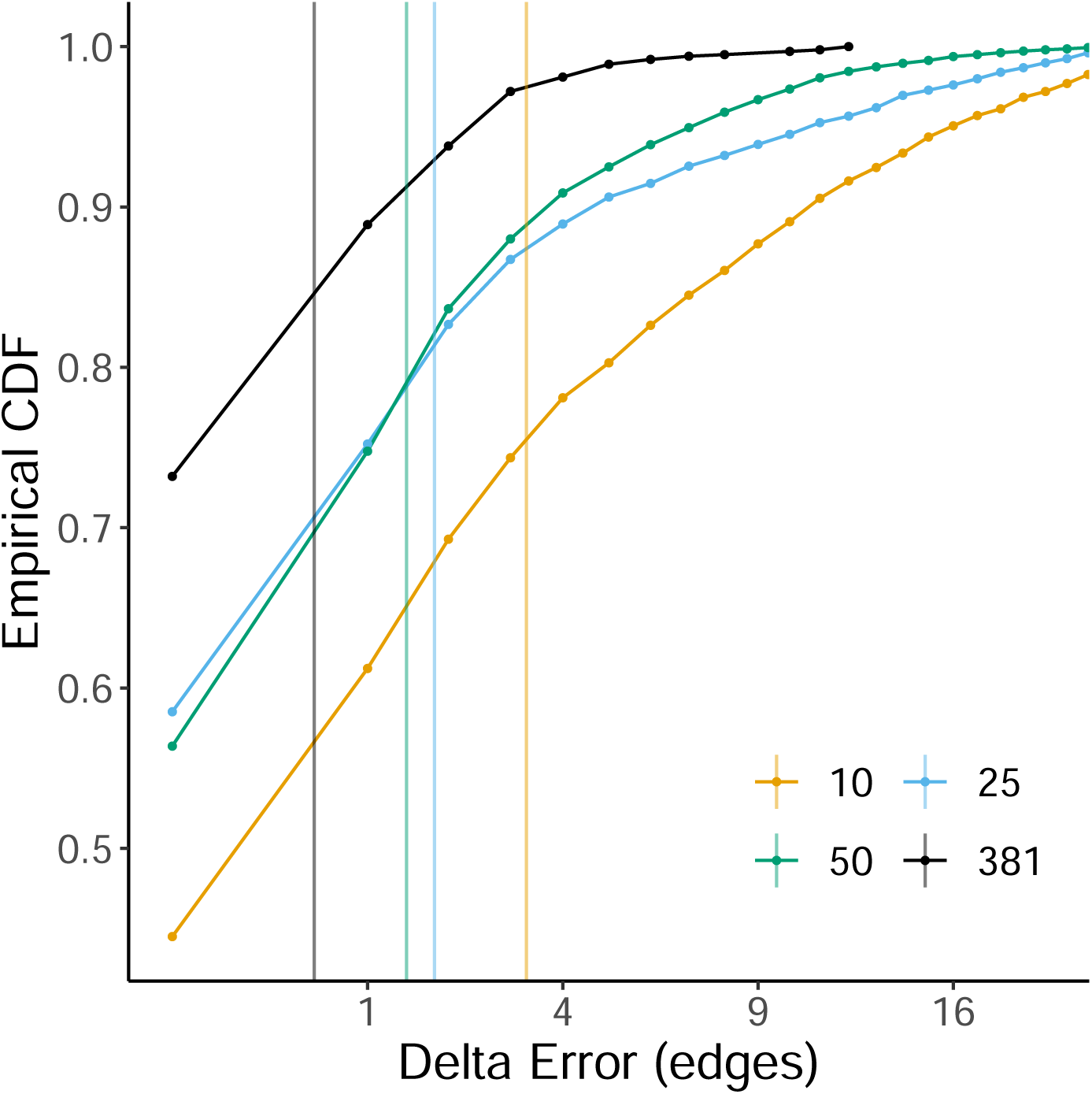
Placement accuracy vs size of the marker gene set coupled with random selection strategy. As the number of genes increases, average error decreases consistently.

**Figure S6:**
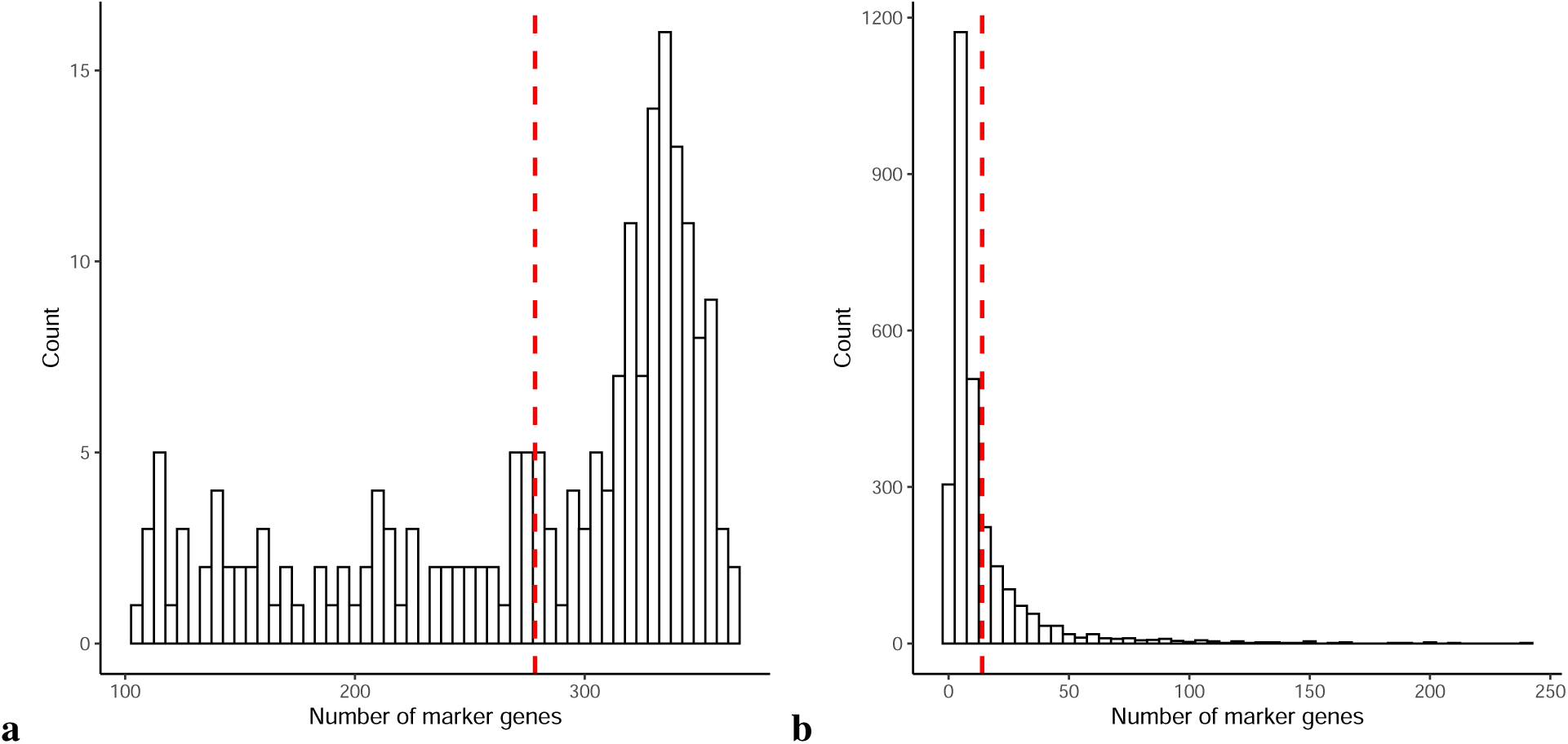
Distribution of the number of marker genes in assemblies and scaffolds. Red dashed line shows the mean. Left: assemblies. Right: scaffolds. Note that most scaffolds have relatively few genes.

**Figure S7:**
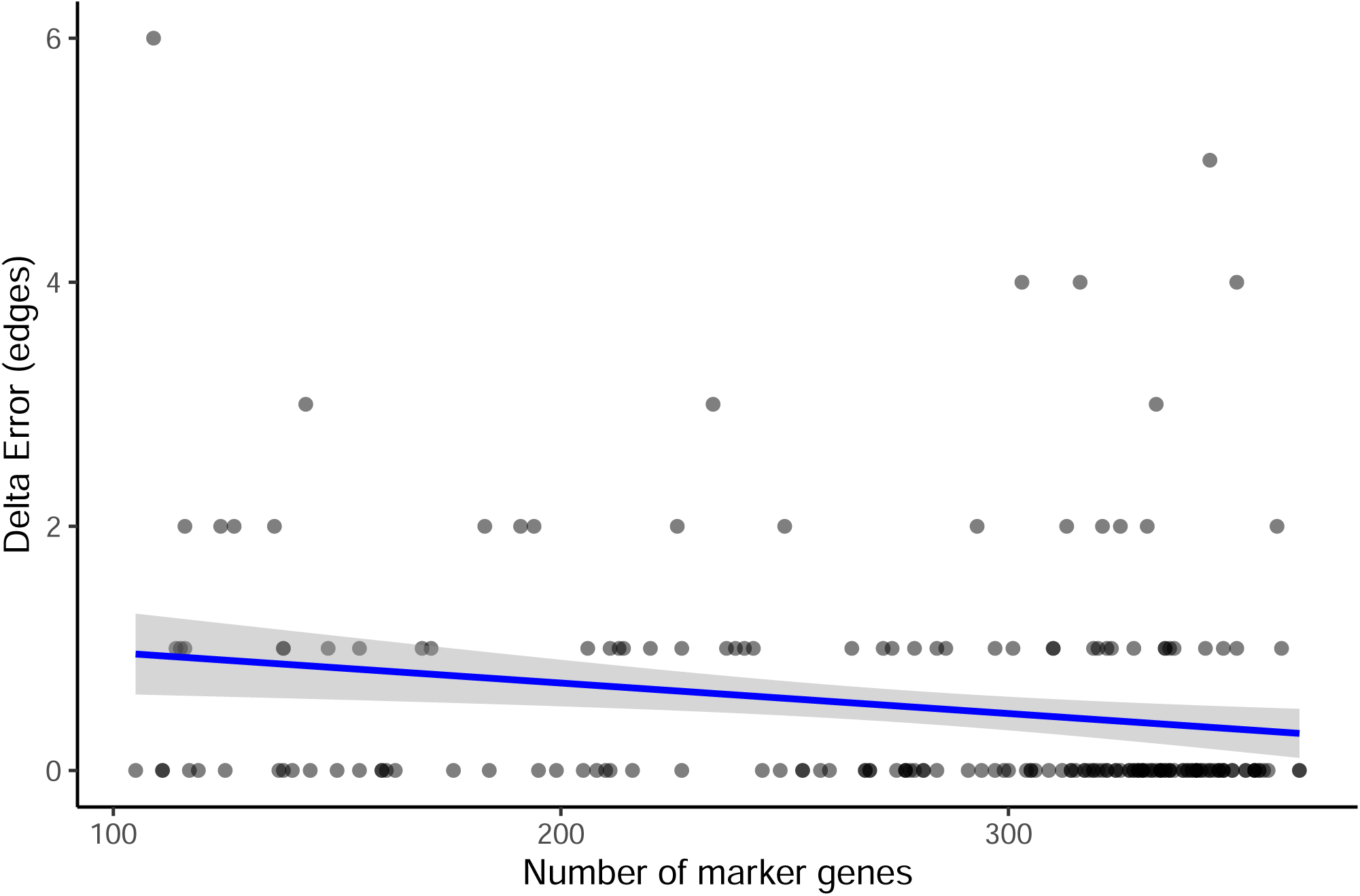
Number of marker genes vs placement error for assemblies. There is a weak but statistically significant correlation between the number of genes and error for placing assemblies.

**Figure S8:**
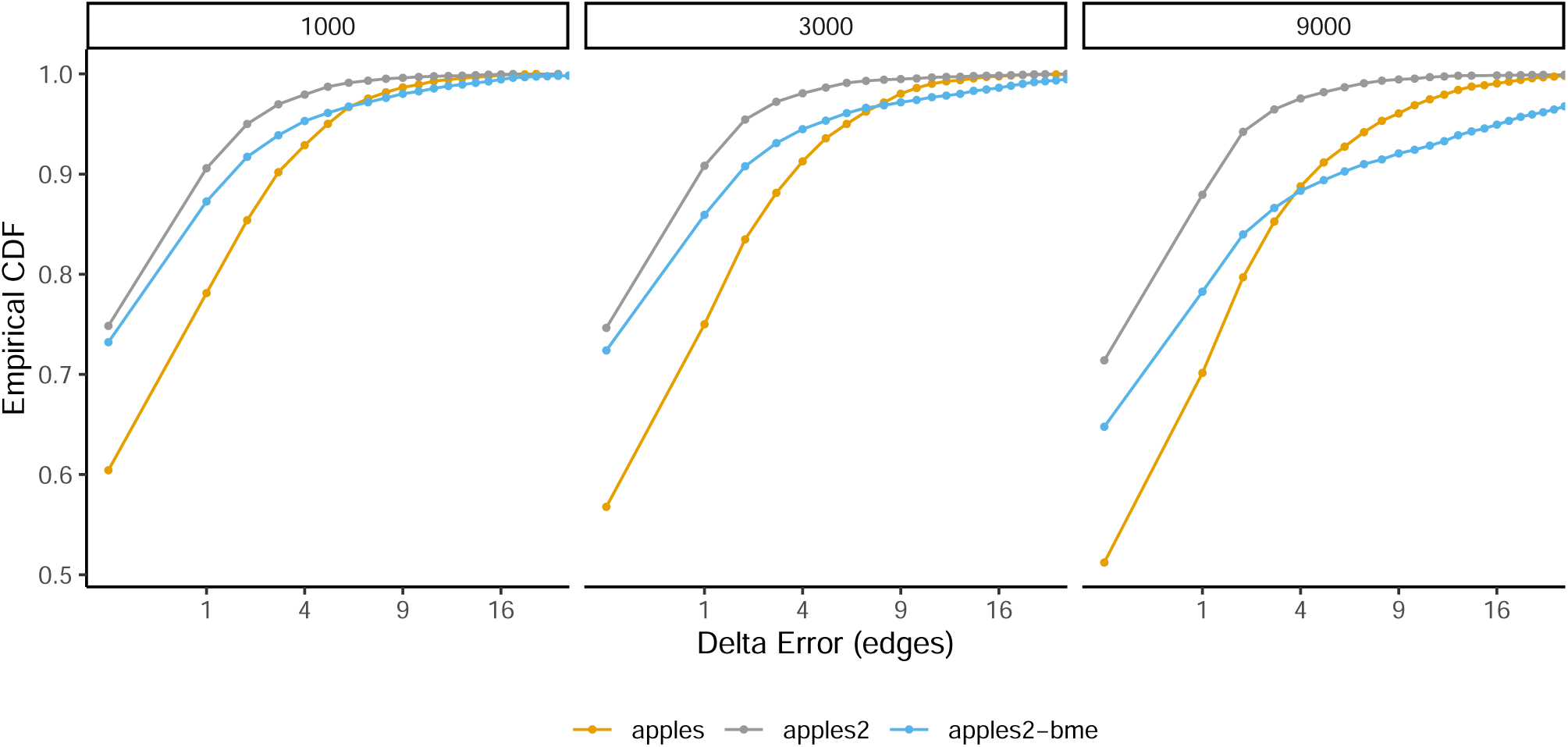
We compare balanced minimum evolution (BME) weighting to Fitch-Margoliash (FM), which is the default weighting scheme in APPLES and APPLES-2.

